# Fitness costs and benefits in response to artificial artesunate selection in *Plasmodium*

**DOI:** 10.1101/2022.01.28.478164

**Authors:** Manon Villa, Arnaud Berthomieu, Ana Rivero

## Abstract

Drug resistance is a major issue in the control of malaria. Mutations linked to drug resistance often target key metabolic pathways and are therefore expected to be associated with biological costs. The spread of drug resistance depends on the balance between the benefits that these mutations provide in the drug-treated host and the costs they incur in the untreated host. The latter may therefore be expressed both in the vertebrate host and in the vector. Research on the costs of drug resistance focusses on interactions with vertebrate host, yet whether they are also expressed in the vector has been overlooked. In this study, we aim to identify the costs and benefits of resistance against artesunate (AS), one of the main artemisinin derivatives used in malaria-endemic countries. For this purpose, we compared different AS-selected lines of the avian malaria parasite *Plasmodium relictum* to their ancestral (unselected) counterpart. We tested their within host dynamics and virulence both in the vertebrate host and in its natural vector, the mosquito *Culex quinquefasciatus*. The within-host dynamics of the AS-selected lines in the treated birds was consistent with the phenotype of resistance described in human *P. falciparum* malaria: a clearance delay during the treatment followed by a recrudescence once the treatment was interrupted. In the absence of treatment, however, we found no significant costs of resistance in the bird. The results of the two experiments to establish the infectivity of the lines to mosquitoes point towards a decreased infectivity of the drug-selected lines as compared to the ancestral, reference one. We discuss the potential implication of these results on the spread of artesunate resistance in the field.

## Introduction

Malaria is a life-threatening disease caused by *Plasmodium* parasites which are transmitted to people through the bites of infected mosquitoes. According to the latest World Health Organization Malaria Report, in 2020 there were an estimated 240 million episodes of malaria which caused over 500 000 deaths, most of them among African children (WHO 2021).

Currently, four major antimalarial drug classes exist for the treatment of malaria: quinolines, antifolates, atovaquone and artemisinin derivatives. The widespread use of these synthetic anti-malarials has saved thousands of lives, but has also exerted an intense selection pressure for the evolution of resistance in the parasite. Drug resistance in malaria has become a major human health issue because *Plasmodium* parasites have evolved resistance to all classes of anti-malarials that have gone into widespread use. The rapid evolution of drug resistance in *Plasmodium* highlights the urgent need to understand the selective pressures under which drug resistant mutations emerge and spread in the population (Read and Huijben 2009; Huijben et al. 2011).

The fate of mutations responsible for resistance is expected to be governed by the balance between the benefits accrued in the drug-treated hosts and the costs suffered in the untreated hosts. The benefits of resistance are obvious: drug resistant parasites show an improved ability to survive the drug treatment. The costs of resistance, on the other hand, arise from the metabolic costs of detoxification or the reduced biochemical efficiency associated with target site mutations and are expected to become manifest in the absence of the drugs (Sirawaraporn et al. 1997; Hastings and Donnelly 2005). For this reason, these costs can be expressed both in the vertebrate host, but also in the vector. Current views about the impact of drug resistance on parasite fitness are, however, almost entirely based on data obtained from vertebrate hosts. In humans, costs of resistance have been inferred by the decrease in the frequency of drugresistance alleles when drug use is stopped or discontinued (Laufer and Plowe 2004; Ord et al. 2007; Babiker 2009). The largest contributions to our understanding of the costs of drug resistance in the vertebrate host have, however, come from work carried out in animal models. *In vivo* competition experiments using laboratory selected drug resistant strains of the rodent malaria parasite *Plasmodium chabaudi* have shown that drug resistant strains are competitively suppressed by the sensitive strains in the absence of drug treatment (De Roode et al. 2004; Wargo et al. 2007; Huijben et al. 2010).

The costs of resistance in the mosquito vector have, in contrast, either been ignored entirely or given only cursory attention (Koella 1998). Yet, the passage through a mosquito is an essential step in the life cycle of *Plasmodium*. Within mosquitoes the parasites reproduce sexually, differentiate, proliferate and migrate to the salivary glands to ensure transmission to the next host. Costs of drug resistance may be expressed at any one of these stages, profoundly impacting the probability of transmission of drug resistant strains. These costs may be expressed in two different ways: by altering the ability of the parasite to infect the mosquito vector (infectivity), or by altering the effects that the parasite has on the fitness of the mosquito vector (virulence).

Here, we investigate the costs and benefits of resistance to artesunate (one of the main artemisinin derivatives) both in the vertebrate host and in the vector. Artemisinin derivatives are the most potent and effective anti-malarial to date. The precise mode of action of artemisinin derivatives is still controversial, but several lines of evidence indicate that they exert their anti-malarial action by perturbing redox homeostasis and haematin detoxification in the parasite (O’Neill et al. 2010). To prevent the evolution of resistance, the World Health Organisation recommends the use of the so-called artemisinin combination therapies (or ACTs) which combine the highly toxic, but short lived, artemisinin derivatives with a partner drug with a long half-life such as sulfadoxine-pyrimethamine, amodiaquine, or lumefantrine (WHO 2021). Despite the hopes raised by the implementation of ACTs, artemisinin resistance is already present in several countries of the Greater Mekong Subregion (WHO 2021), and worryingly, the *de novo* emergence of artemisinin resistance has been recently detected in Subsaharan Africa (Uwimana et al. 2020). Phenotypically, artemisinin resistant parasites are characterized by (i) a delay in parasite clearance (resistant parasites take an extra 24-48 to be cleared from the blood) followed by (ii) a recrudescence of the parasite at the end of the standard 3-day therapeutic course (Dondorp et al. 2009). This phenotype has been shown to be associated to several mutations in the Kelch 13 (K13) propeller domain of the parasite (Straimer et al. 2014). Despite the ongoing spread of artemisinin resistance across malaria-endemic areas *we* still lack a clear picture about the costs and benefits associated to these mutations (but see (Nair et al. 2018; Tirrell et al. 2019).

Selecting for drug resistance in rodent and avian malaria has proven to be a straightforward procedure that provides a powerful tool for testing the effects of a particular drug resistance mutation on parasite fitness *in vivo* (Greenberg 1956; Beaudoin et al. 1967; De Roode et al. 2005; Walliker et al. 2005; Wargo et al. 2007; Huijben et al. 2011). Here we select for artesunate (AS, a widely-used artemisinin derivative) resistance in *Plasmodium relictum* (the most widespread species of avian malaria in the wild) by submitting it to gradually increasing doses of the drug. We then compare the within host dynamics and virulence of the drug-selected lines to the reference line both in the presence and in the absence of an AS treatment in the vertebrate host. We expect that: i) in treated hosts (birds), artesunate-selected lines will exhibit a similar phenotype to human malaria, i.e. delayed clearance followed by recrudescence, ii) in untreated hosts (birds) and vectors (mosquitoes), artesunate-selected parasites suffer higher fitness costs in comparison to their unselected counterparts.

## Methods

### Mosquito and parasite protocols

*Plasmodium relictum* (lineage SGS1) is the aetiological agent of the most prevalent form of avian malaria in Europe. The biology of this parasite species is similar to that of human malaria, both in the vertebrate host and in the mosquito. Avian malaria has historically played a key role in human malaria research, and in particular in the development and testing of the first anti-malarials (Hewitt 1940; Rivero and Gandon 2018). Our parasite lineage was isolated from blue tits (*Parus caeruleous*) collected in the Montpellier area (France) in October 2016 and subsequently passaged to naïve canaries (*Serinus canaria*)by intraperitoneal injection. Since then, it has been maintained by carrying out regular passages between our stock canaries through intraperitoneal injections with the occasional passage through the mosquito (one passage through mosquito every c.a. 15 bird to bird passages, on average).

All experiments were carried out using a laboratory line of *Culex quinquefasciatus* (SLAB strain). Vectors in the *Culex pipiens* complex (*Cx pipiens* and *Cx quinquefasciatus*) are the principal natural vector of *Plasmodium relictum* in the wild. The larvae in all the experiments were reared at a constant density per tray (n=300 larvae) following previously published laboratory protocols (Vézilier et al 2010). Larval trays (n=20) were placed individually inside an “emergence cage” (40 cm x 28 cm x 31 cm) and emerged adults were allowed to feed *ad libitum* on a 10% glucose water solution.

AS-selected parasite lines were obtained by treating infected birds with increasing concentrations of artesunate (Sigma A3731). Birds were infected by injecting them intra-peritoneally with 100 μl of infected blood. The artesunate treatment was initiated 12 days later, to coincide with the peak of the acute phase of the parasite’s infection. Artesunate was diluted in bicarbonate (50 mg/Kg) and administered through intra-peritoneal injections carried out twice daily (9am and 6pm) for four consecutive days. Five or six days later, 100 μl of blood from the treated birds was taken via a wing puncture, transferred to new canaries, and the treatment was thus repeated until a total of 5 passages were carried out. Three AS-selected lines were obtained in this way. Line AS1 was obtained by treating birds with 16 mg/kg artesunate for 3 passages followed by 32 mg/kg for a further 2 passages. Lines AS2 and AS3 were obtained by treating birds with 8 mg/kg for 3 passages followed by 16 mg for 2 passages. The reference line was maintained in parallel by injecting birds with bicarbonate following the same protocol. Experiments were initiated immediately after the 5^th^ passage.

### Experiment 1. Phenotypic characterization of drug resistance in the bird

Artesunate-resistance is expected to be beneficial for the parasite in the presence of the drug, but to incur fitness costs when the drug is absent. In this experiment, we infected birds with either the reference line or one of the drug-selected lines (AS1, AS2, AS3). Half of the birds where then treated with a standard dose of artesunate, while the other half were given a sham (bicarbonate) injection. For each bird we quantified parasitaemia (proportion of red blood cells infected) through time and parasite virulence (bird weight loss and anaemia).

The detailed experimental protocol was as follows (**Figure 1**). Birds were inoculated with each of the parasite lines (reference, AS1, AS2 and AS3) by means of an intra-peritoneal injection (6 birds per line, all birds in these experiments were 1-2 years old). Each bird received between 70 000 to 90 000 parasites, except for the birds injected with the AS3 line, which received 30 000 parasites because the parasitaemia in the donor birds were too low. Bird weight and red blood cell count (RBC: number of red blood cells per ml of blood, Beckman Coulter Counter, Series Z1) were measured immediately before the infection and every two days thereafter. Bird parasitaemia (% red blood cells infected) was monitored every two days from the second day after the inoculation (henceforth “day 2 pbi”: post-bird inoculation) onwards, using thin blood smears. On day 12 pbi, to coincide with the acute phase of the parasite infection, half of the birds were injected with 50 μl of 16mg/kg AS (dissolved in 50mg/kg bicarbonate) while the other half were injected with an equivalent amount of bicarbonate (50mg/kg). The injections took place twice a day (9 am and 6 pm) for 4 consecutive days (days 12-15 pbi). During this period, parasitaemia and virulence measurements were taken 24h after each injection (days 13-16 pbi). Bird parasitaemia, weight and RBC were thereafter followed every two days for 16 days after the end of the treatment (days 18-30 pbi, **Figure 1**).

**Figure 1:**
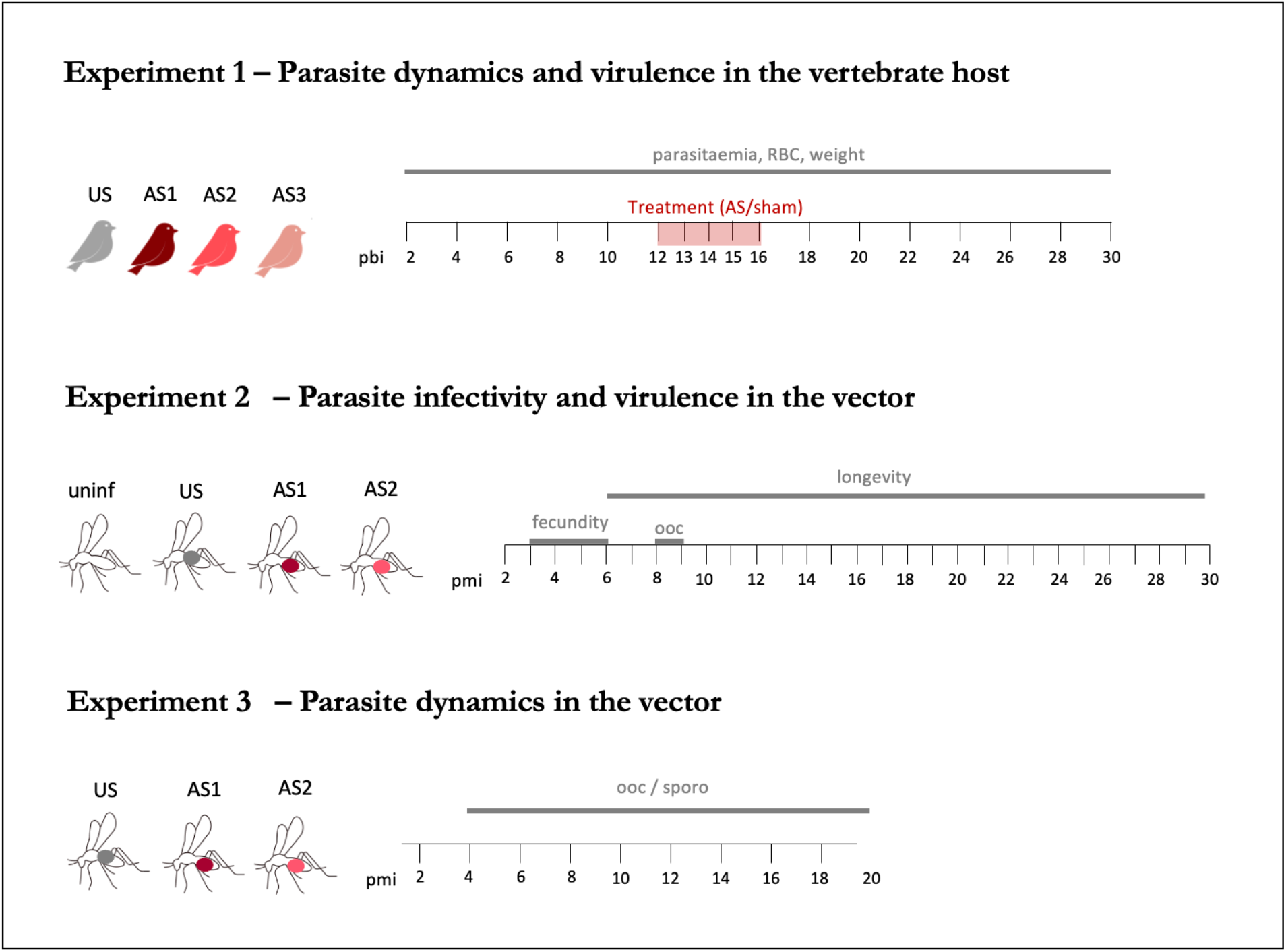
Experimental design for experiments 1 - 3. In Experiment 1 (n= 6 birds per line), samples were taken every two days between day 2 and day 30 post bird inoculation (pbi), except during the treatment course where samples were taken daily. The parameters quantified were: bird parasitaemia, bird weight and RBC count. In Experiment 2 (n = 2 birds per line plus 2 uninfecteed birds) we quantified fecundity (4-7 pmi, ~30 mosquitoes per bird per line), longevity (~150 mosquitoes per bird per line, survival was monitored on a daily basis), and oocyst count (8-9 pmi, 20 mosquitoes per bird). In Experiment 3 (n = 3 birds per line): we dissected mosquitoes every two days (2-20 pmi) and quantified oocysts (microscopic examination of the midgut) and sporozoites (qPCR of the thorax). pbi = days post bird infection, pmi = days post mosquito infection. US = unselected reference line, AS1, AS2, AS3 = artesunate-treated (putatively resistant) lines.

To account for the variance in parasitaemia between birds on the day immediately preceding the treatment (day 12 pbi, see Supplementary Table ST1), the parasitaemia on day 13 pbi onwards was calculated as a fold-increase (or decrease) in the number of parasites using the parasitaemia at 12 pbi as the baseline (*px/pb*, where *px* is the parasitaemia at a given day, and *pb* the baseline parasitaemia at day 12 pbi).

### Experiment 2. Costs of drug resistance in the vector: parasite infectivity and virulence

The aim of this experiment was to establish whether AS-selected and reference lines differ in their infectivity and virulence to the mosquito. For this purpose, mosquitoes were allowed to blood feed on birds infected with the drug-selected and reference parasite lines. Mosquitoes fed on uninfected birds were used as controls. A subsample of the mosquitoes was used to quantify parasite density, while the rest were followed for several days to measure their fecundity and longevity. Since having too many mosquitoes in the cage risk compromising the wellbeing of the bird, and given that AS3 did not have the expected phenotype of resistance (delayed clearance followed by recrudescence), this experiment was done using only AS1 and AS2.

The detailed experimental protocol is as follows (**Figure 1**). Six naïve birds were infected with 100 000 parasites of either the reference, AS1 or AS2 lines (2 birds per line). Two further birds were used as uninfected controls. On day 12 pbi, birds were introduced individually into cages containing 150 12-day old female mosquitoes. The following morning, unfed females were counted and discarded from the study and all cages were provided with *ad libitum* sugar in the form of a 10% sugar solution. To quantify fecundity, three days after the blood meal (day 3 pmi, post-mosquito infection), a tray of water was placed in each to allow females to lay eggs. The trays were checked daily for the presence of eggs for 3 consecutive days (days 3-6 pmi). *Culex pipiens* females lay eggs in a single raft containing up to 250 tightly-packed eggs. This facilitates the estimation of the number of laying females and the eggs laid per female. The egg laying date was recorded and the egg rafts were photographed using a binocular microscope equipped with a numeric camera. The eggs counted using the Mesurim Pro freeware (Academie d’Amiens, France). Survival was assessed daily by counting dead individuals lying at the bottom of each cage until all females died.

To quantify parasitaemia, on day 8 and 9 pmi, 20 mosquitoes were haphazardly sampled from each cage and immediately dissected under a binocular microscope to extract the oocyst-infected midguts. Midguts were stained with a 5% mercurochrome solution to assess infection rate (oocysts present/absent) and oocyst burden (number of oocysts) under a phase contrast microscope.

### Experiment 3. Costs of drug resistance in the vector: parasite dynamics

The aim of this experiment was to explore whether the results obtained in Experiment 2 may be explained by a difference in the within-vector parasite dynamics between AS-selected and reference lines. For this purpose, mosquitoes were allowed to blood feed on birds infected with the drug-selected and reference parasite lines (as above). Mosquitoes were dissected at regular intervals and the number of oocysts and sporozoites were compared between the AS-selected and the unselected reference lines.

The detailed experimental protocol is as follows (see **Figure 1**). Six naïve birds were infected with 100 000 parasites of either the reference, AS1 or AS2 lines (2 birds per line). As in the previous experiment, on day 12 pbi, birds were introduced individually into cages containing 150 12-day old female mosquitoes. The following morning, unfed females were counted and discarded from the study and all cages were provided with *ad libitum* sugar for the reminder of the experiment. Starting on day 4 post blood meal and every 2 days thereafter (4-20 dpmi), 20 mosquitoes were haphazardly sampled from each cage and immediately dissected under a binocular microscope to extract the oocyst-infected midguts and count the oocysts (as above). The mosquito thorax, containing the sporozoite-infected salivary glands, was then severed and frozen at −20°C for the subsequent quantification of parasites via qPCR. Sporozoite quantification was calculated as a ratio between mosquito DNA (*ace2* gene) and *Plasmodium relictum* DNA (*cytb*) as described previously (Zélé et al. 2014).

### Ethics statement

Bird manipulations were carried out in strict accordance with the “National Charter on the Ethics of Animal Experimentation” of the French Government. Experiments were approved by the Ethical Committee for Animal Experimentation established by the authors’ institution (CNRS) under the auspices of the French Ministry of Education and Research (permit number CEEA-LR-1051).

### Statistical analyses

Analyses were carried out using the R statistical package (v3.4.4). The general procedure to build models was as follows: line (reference, AS1, AS2, AS3), sampling day, treatment (AS treated or sham-treated, Experiment 1 only) and parasitaemia were introduced into the model as fixed explanatory variables. Bird ID and the qPCR plates used to quantify sporozoites in the head-thorax fraction of the mosquito (Experiment 3 only) were fitted as random effects. Maximal models, including all higher order interactions, were simplified by sequentially eliminating non-significant terms and interactions to establish a minimal model. The significance of the explanatory variables was established using a likelihood ratio test (LRT) which is approximately distributed as a chi-square distribution (Bolker 2008) and using p = 0.05 as a cut-off p-value.

Mosquito survival (Experiment 2) was analyzed using Cox proportional hazards mixed effect models (coxme). Proportion data (oocyst and sporozoite prevalence) were analyzed using mixed linear models and a binomial distribution. Count data (eggs per raft) were analyzed using mixed linear models and a Poisson distribution. Response variables that were highly overdispersed (e.g. oocyst burden) were analyzed using mixed negative binomial models (glmmTMB). *A posteriori* contrasts were carried out by aggregating factor levels together and by testing the fit of the simplified model using a LRT (Crawley 2007). Statistical analyses are summarised in Supplementary Table ST2. Two additional standard measurements of survival were obtained in each cage: the median survival (the time at which 50% of the population is still alive) and the proportion of mosquitoes that survived till day 14 (the time at which sporozoites peak in all lines, see below).

## Results

### Experiment 1. Phenotypic characterization of artesunate resistance in bird

We infected birds with either the reference line or one of the drug-selected parasite lines (AS1, AS2, AS3). Twelve days later, half of the birds were then treated with AS, while the other half were given a control (bicarbonate) injection (**Figure 1**). For each bird, we quantified parasite density dynamics (proportion of infected red blood cell through time) and parasite virulence (bird weight loss and anaemia) before, during and after the AS treatment.

We started by analysing whether there were differences in parasitaemia between lines *before* the AS-treatment (days 2-12 pbi). We fitted a full model that included day, line and treatment (and bird as a random factor to control for temporal pseudo-replication). There was no significant 3-way interaction and no significant differences in parasitaemia between birds infected with the different lines, or subsequently allocated to the treated or control treatments (although variance between birds was high, **Supplementary Figures SF1A-D**). This model only detected differences in parasitaemia between days (model 1, χ^2^= 33.34, p < 0.0001).

*During* the treatment (days 13-16 pbi), the parasitaemia of treated and untreated birds followed very different courses. Treatment with AS had a very significant effect on the parasitaemia, and the effect was dependent on both the day and the parasite line (model 2, χ^2^= 12.088, p = 0.0071, **Figure 2A**). To obtain a more accurate picture of the nature of this interaction, subsequent analyses were done separately for AS-treated and reference birds. Treatment with AS resulted in an overall decrease in parasitaemia in all lines (see Supplementary Table ST3). On the first day after the treatment (day 13 pbi) the reduction in parasitaemia was greater in the reference line (− 93.6% on average) than in either the selected AS1 (−59.8%), AS2 (− 57.8) or AS3 (− 69,6%) lines. This effect largely disappears in subsequent days (Supplementary Table ST3). The statistical analyses revealed a significant interaction between day and line (model 3, χ^2^= 20.563, p= 0.0147). In untreated birds, there was a significant difference between the lines (model 4, χ^2^= 9.1288, p= 0.0276) due to AS2 having a significantly higher parasitaemia than the other lines. This effect was independent, albeit marginally, of the sampling time (model 4, χ^2^= 7.7845, p= 0.0507, **Figure 2B**).

**Figure 2:**
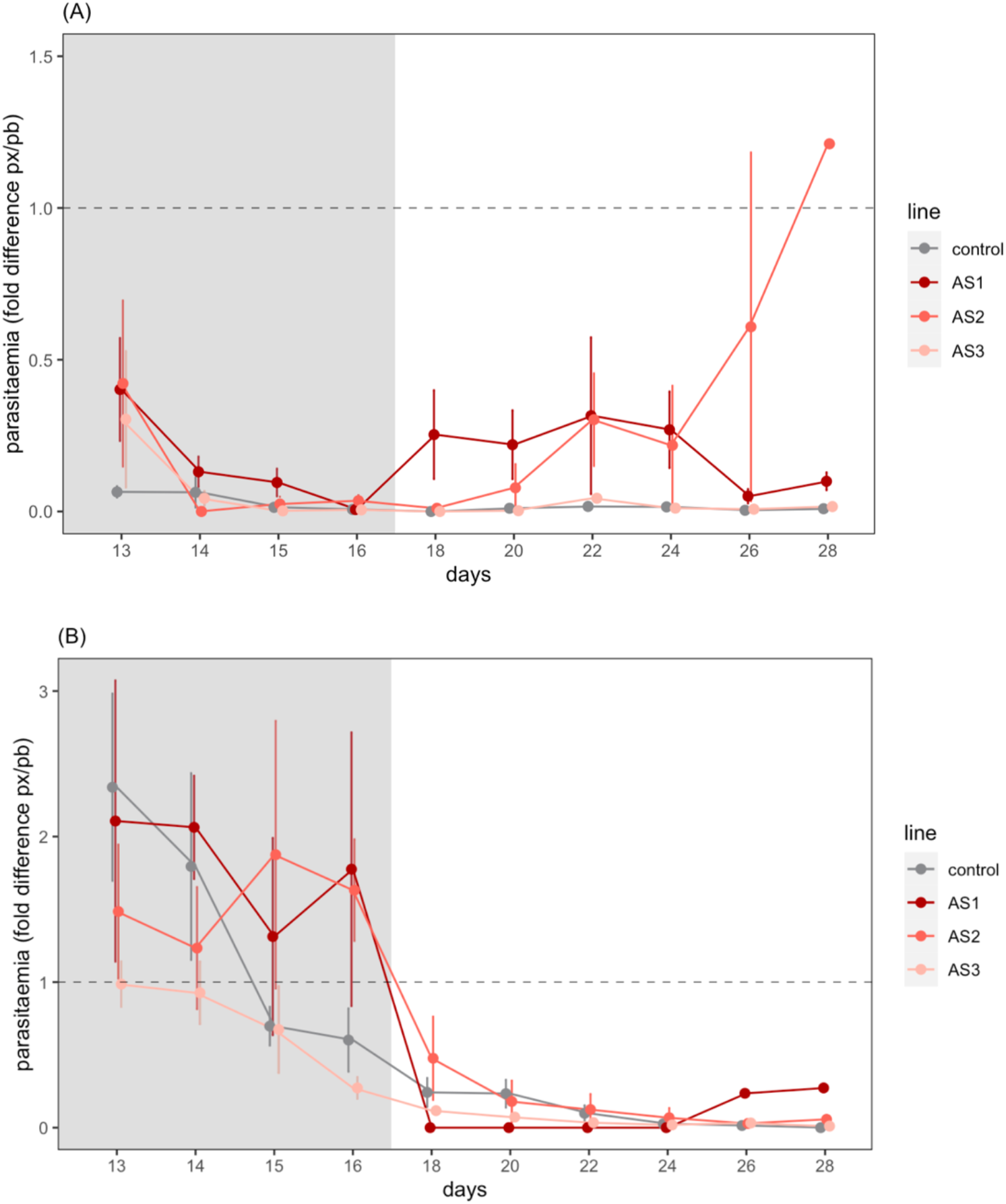
Experiment 1. Parasitaemia (mean ± s.e) in artesunate-treated (**A**) and untreated (**B**) birds. Shaded area corresponds to the 4-day treatment period (measurements taken 24h after each injection). Untreated birds were sham injected with the artesunate solvent. Dashed line indicates the baseline parasitaemia on the day immediately before the first injection (day 12). Bars above/below the dashed line indicate an increase/decrease in parasitaemia with respect to day 12.

*After* the AS treatment, parasitaemia in the treated birds showed a different pattern between the lines (model 5, χ^2^= 8.8851, p= 0.0308, **Figure 2A**). Lines AS1 and AS2 had a significantly higher parasitaemia than the reference line. In untreated birds, the parasitaemia decreased gradually with time (model 8, χ^2^= 54.449, p < 0.01) and was independent of the parasite line (model 6, *line*: χ^2^= 7.9084, p= 0.0479, **Figure 2B**).

Virulence was quantified as a change in weight and a change in RBC counts (anaemia) with respect to the weight and RBC counts on the first day of the experiment. Weight changes following the infection were significant and dependent both on the parasite line, on the treatment group (AS-treated or untreated), and on the timing with respect to the treatment (before, during or after, model 7, χ^2^ = 34.51, p = 0.001). In untreated birds, there was a weight difference between the lines, with birds infected with AS1 and AS2 (but not AS3) having a significant lower weight than those infected with the reference line throughout the experiment (model 9, χ^2^ = 26.56, p = 0.0002, Figure SF2B). In treated birds, however, the weight difference disappeared during the treatment, only to reappear after the treatment ended (model 8, χ^2^ = 29.28, p < 0.0001, **Supplementary Figure SF2A**). RBC counts were significantly lower during the treatment than before or after the treatment (model 10, χ^2^ = 54.55, p < 0.0001), but were independent of the line and were similar in AS-treated and control birds (**Supplementary Figures SF3A and SF3B**).

### Experiment 2. Costs of drug resistance in the vector: parasite infectivity and virulence

Mosquitoes were allowed to blood feed either on uninfected birds or on birds infected with drug-selected and reference parasite lines. A subsample of the mosquitoes fed on the infected birds was used to quantify oocyst density, while the rest were followed for several days to quantify their fecundity and longevity.

The overall prevalence of infection was low (29 ± 4% of female mosquitoes had at least 1 oocyst in their midgut) and independent of the parasite lines (model 11, χ^2^= 0.6017, p= 0.7402, **Figure 3A**). Oocyst burden (number of oocysts in infected females) was, however, significantly different between the lines (model 12, χ^2^= 8.0512, p= 0.0179); mosquitoes infected with the reference line had a significantly higher number of oocysts (mean ± s.e: 70 ± 22) than mosquitoes infected with the AS1 (mean ± s.e: 13 ± 5) and AS2 (mean ± s.e: 25 ± 11) lines (**Figure 3B**).

**Figure 3:**
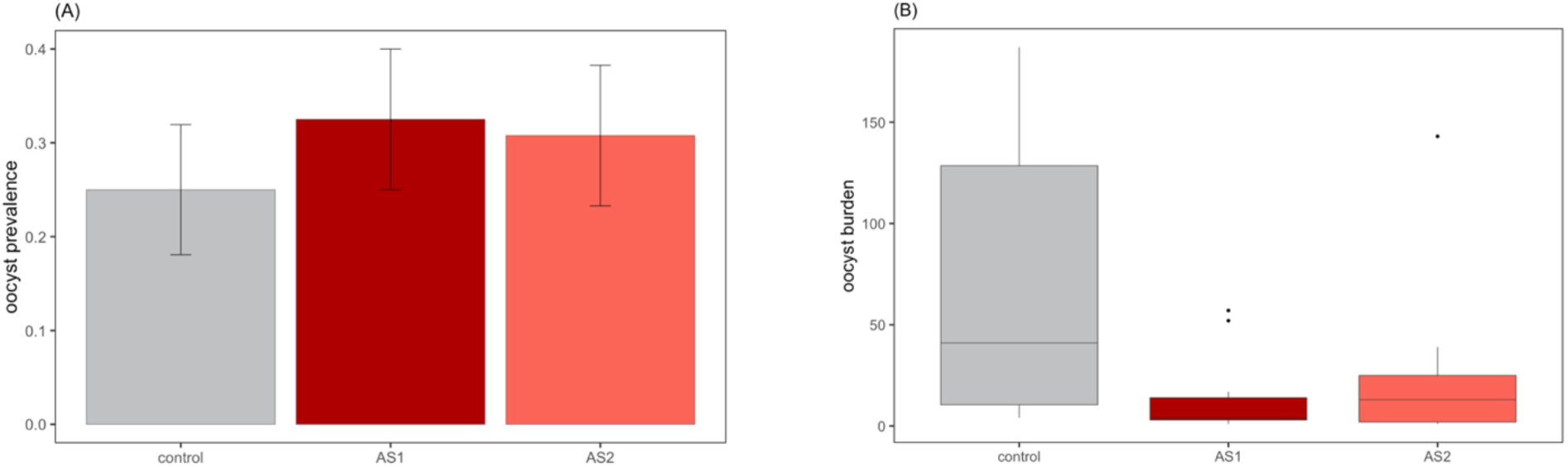
Experiment 2. Mean (± s.e) oocyst prevalence (**A**) and burden (**B**) in mosquitoes infected by each parasite line 8-9 days post infection. Burden is represented as a boxplot showing the median (horizontal lines), and the first and third quartiles (box above and below the medians). Vertical lines delimit 1.5 times the inter-quartile range above which individual counts are considered outliers and marked as circles.

There were also significant differences in mosquito fecundity (number of eggs per raft) between lines (model 14, χ^2^= 11.1, p= 0.0112). Post-hoc analysis showed that mosquitoes infected with the AS2 line laid significantly fewer eggs (mean ± s.e: 126 ± 4) than uninfected mosquitoes (142 ± 5) and mosquitoes infected with the refence (142 ± 4) and AS1 (148 ± 4) lines (χ^2^= 6.5741, p= 0.0104, **Figure 4**). There were no significant differences in longevity between infected and uninfected mosquitoes, or between mosquitoes infected with the different lines (model 13, χ^2^= 3.7758, p= 0.2867). The median longevity (the time at which 50% of the mosquitoes in each treatment is still alive) and the proportion of mosquitoes that survived till day 14 (peak sporozoite production) are given in Supplementary Table ST4.

**Figure 4:**
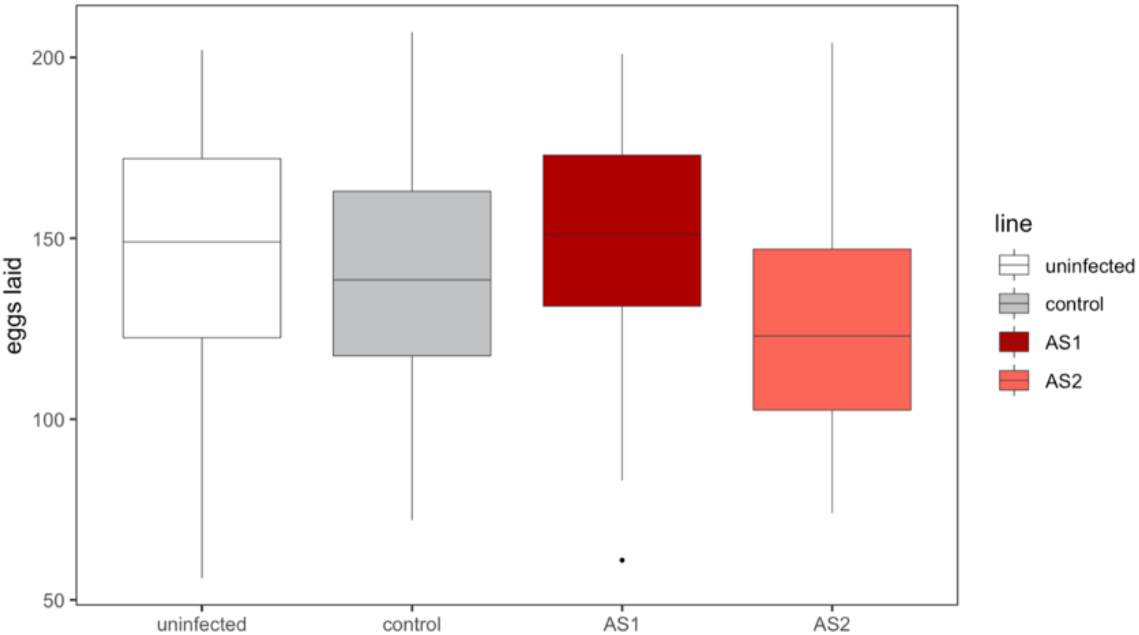
Experiment 2. Fecundity of *Culex pipiens* females uninfected (n=60) or infected with each of the different parasite lines (n=60 per line). The number of eggs per raft is represented as a boxplot with the median (horizontal lines), first and third quartiles (box above and below the medians). Vertical lines delimit 1.5 times the inter-quartile range above which individual counts are considered outliers and marked as circles.

### Experiment 3 Costs of drug resistance in the vector: parasite dynamics

This experiment was carried out to investigate whether the differences in oocyst burden between selected and reference lines may be due to a difference in the dynamics of development of the parasite within mosquitoes. **Figure 5** shows the cumulative increase in oocysts in mosquitoes having fed in birds infected with each of the lines (reference or AS-selected). No mosquitoes in any of the lines were infected on day 4 so this time point was taken off the analyses. Mosquitoes infected with the reference line showed a faster cumulative increase in oocysts than mosquitoes infected with the AS-selected ones. As a result, by days 8-9 (corresponding to the dissection time in Experiment 2), mosquitoes fed on birds infected with the reference line had between 60-70% of the cumulative total number of oocysts, while mosquitoes fed from 3 out of the 4 birds infected with AS-selected lines had 0-16% of the cumulative total. Mosquitoes fed from one of the (AS1-infected) bird, however, showed an unusually high number of oocysts on the first dissection day.

**Figure 5:**
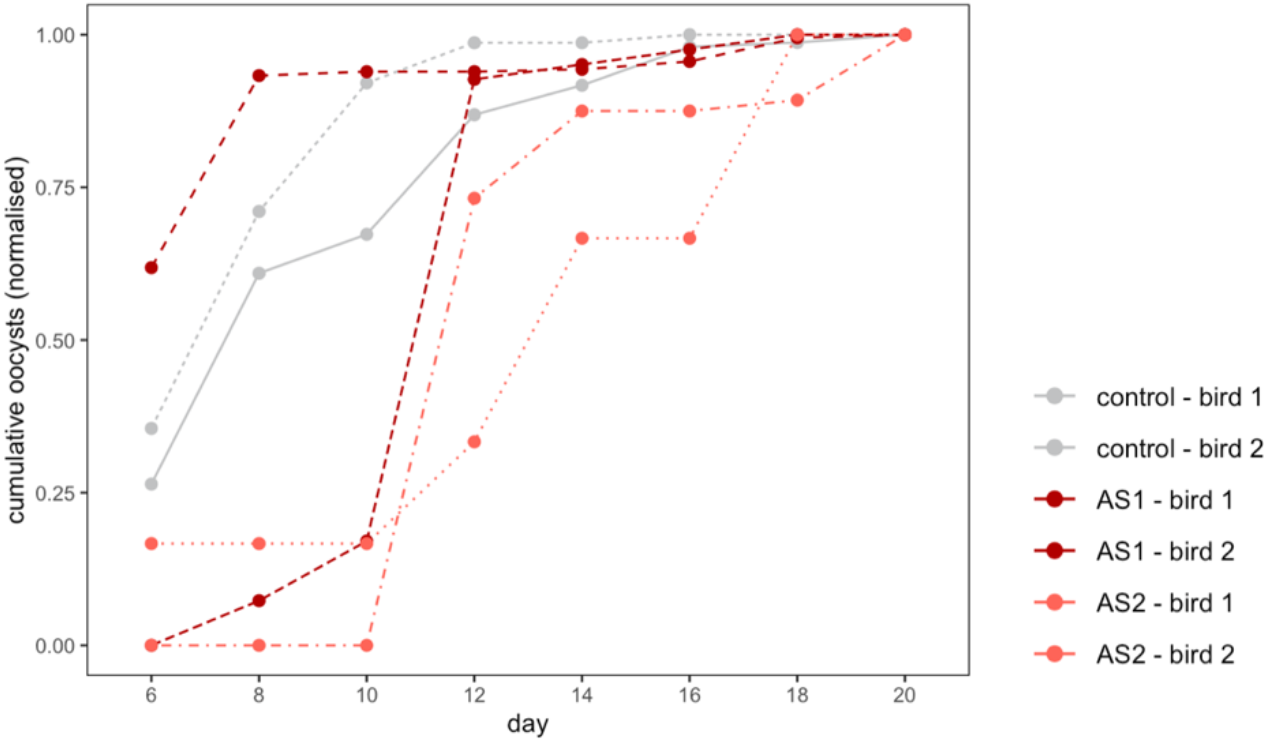
Experiment 3. Cumulative proportion of oocysts per day in mosquitoes infected with the reference (control) and AS-selected lines. Data is provided separately for each cage/bird.

The mean oocyst prevalence (% mosquitoes containing at least 1 oocyst) averaged over the whole experiment was higher in reference (67.92 ± 6.47%, mean ± s.e.) and AS1 lines (53.65 ± 7.88%) than in the AS2 line (36.36 ± 10.49%). The same pattern was found with oocyst burden, which was higher in the reference (32.80 ± 11.95) and AS1 lines (18.90 ± 11.33) than in the AS2 line (7.75 ± 4.83). When day was fitted to the model the effect of line, however, disappeared for both oocyst prevalence (model 15, χ^2^= 5.3164, p= 0.070 and burden (model 16, χ^2^= 0.2313, p= 0.8908; **Figure 6A and 6B**).

**Figure 6.**
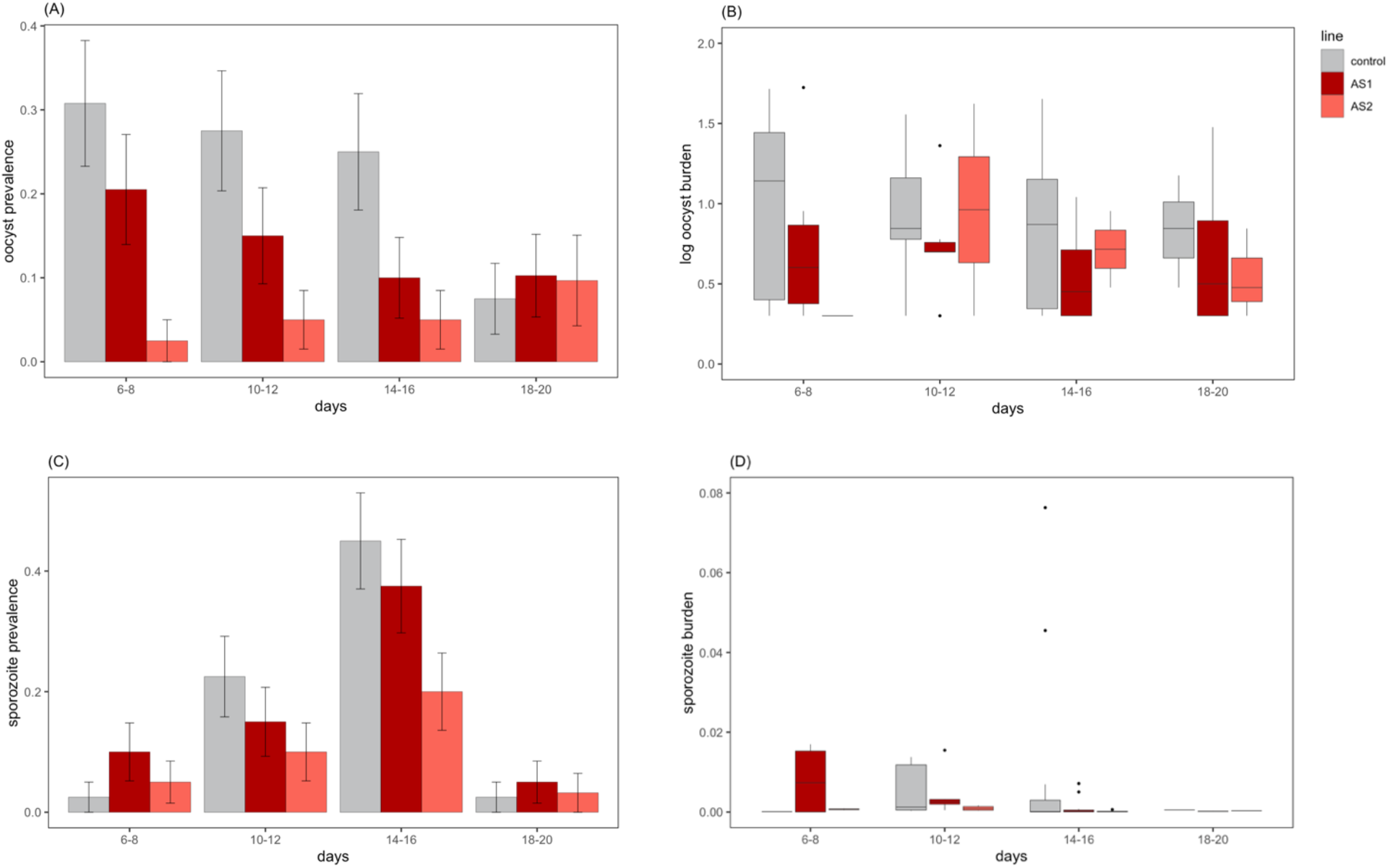
Experiment 3. Oocyst prevalence (A) and burden (B), sporozoite prevalence (C), and burden (D) of mosquitoes infected by each of the parasite lines for different sampling intervals. Prevalence is represented as the mean ± standard error (calculated as sqrt(pq/n)). Oocyst and sporozoite burden are represented as a boxplot with the median (horizontal lines), first and third quartiles (box above and below the medians). Vertical lines delimit 1.5 times the inter-quartile range above which individual counts are considered outliers and marked as circles.

The mean sporozoite prevalence, averaged over the whole experiment, was roughly 2-times higher in the reference (28.30 ± 6.24%) and AS1 lines (27.90 ± 6.92%) than in the AS2 line (13.63 ± 7.48%). Sporozoite burden, on the other hand, was 4-5 times higher in the reference line (0.0126 ± 0.0067) than in the AS-selected lines (AS1: 0.0026 ± 0.0009, AS2: 0.0034 ± 0.0029). As above, when day was fitted to the model the effect of line on sporozoite prevalence and burden disappeared (model 17, χ^2^= 0.1788, p= 0.9145, and model 18, dχ^2^= 1.9214, p= 0.3826, respectively **Figure 6C and 6D**).

## Discussion

The two telltale signs of *P. falciparum* resistance to artemisinin derivatives in the field are a delay in the clearance of parasites during ACT treatment followed by a recrudescence once the treatment is interrupted (Douglas et al. 2021). Artemisinin derivatives are remarkably efficient at clearing susceptible parasites from the blood (typically within 24-48 hours in *P. falciparum*). A slower clearance of parasites after the treatment, typically taking an extra 24-48 hours, is interpreted as evidence of resistance to the artemisinin-derivative component (Witkowski et al. 2013; Mok et al. 2015). This delayed clearance phenotype is a far cry from the standard definition of resistance (survival to a drug treatment), which is why artesunate resistance is commonly referred to as a ‘partial resistance’ (Uhlemann et al. 2010; Menard and Dondorp 2017). Partial artemisinin resistance rarely leads to ACT failure, but may select for drug resistance to the partner drug, as larger parasite populations remain in the blood stream for longer periods of time (Fairhurst 2015). In our study, the within-host dynamics of the drug-selected lines in the treated birds was consistent with the phenotype of resistance on *P. falciparum* in the field. AS-selected lines took longer to respond to the treatment than their reference line counterparts. In the first 24h after the beginning of the treatment the parasitaemia of the reference line dropped by over 90%, while that of the AS-selected lines only dropped by between 60% (AS1, AS2) and 70% (AS3). By the following day, there was no discernible difference between the parasitaemia of the reference and drug selected lines, (**Figure 2A**). There are no significant differences in the pre-treatment dynamics between the lines that may explain this significant difference in the parasitaemias in the first 24h between the treatments. (**Supplementary Figure SF1**). In keeping with previous studies on *P. falciparum* parasites, two of our drug-selected lines (AS1, AS2) also had a pronounced recrudescence after the treatment was interrupted, producing on average more than 20 times as many parasites in the two weeks following drug treatment than did the reference line (**Figure 2A**). One potential explanation of the different dynamic of AS3, may be found in the lower infective dose given to the birds (see above).

Drug resistance mutations are known to disrupt the parasite’s metabolism, generating fitness costs. In drug-treated hosts these costs are largely compensated by the benefits conferred by the resistance. In untreated hosts, however, the magnitude of these costs will determine whether these mutations will persist and spread in the population. Current views about the impact of drug resistance on parasite fitness are almost entirely based on data obtained from the vertebrate host. In humans, costs of resistance have been inferred by the decrease in drug-resistance alleles when drug use is stopped or discontinued (Laufer and Plowe 2004) and from the decrease in the frequency of drug resistant parasites during the dry season, when there is no transmission and drug use is dramatically reduced. Direct experimental evidence of the costs of artemisinin resistance has been obtained by setting up *in vitro* competition experiments between drug-resistant and reference *Plasmodium* strains (Hott et al. 2015; Nair et al. 2018; Tirrell et al. 2019; Mathieu et al. 2020). To our knowledge the only *in vivo* experiments were carried out with *Plasmodium chabaudi* (a rodent species) and showed no obvious cost of AS in the absence of treatment (Pollitt et al. 2014).

We did not observe any costs associated to the AS-selected lines in the absence of treatment. Experimental evidence of the costs of resistance has been obtained by setting up competition experiments between drug-resistant and susceptible *Plasmodium* strains. Using *in vitro* experiments where the erythrocyte stages of a drug resistant *P. falciparum* strain bearing mutations in the *pfmdr1* gene competed against a drug-sensitive strain with the same genetic background, Hayward et al (2005) estimated that the loss of fitness of the resistant strain was ca. 25% per generation. *In vivo* competition experiments using laboratory selected drug resistant strains of the rodent malaria parasite *Plasmodium chabaudi* have confirmed that drug resistant strains are competitively suppressed by the sensitive strains in the absence of drug treatment (de Roode *et al*. 2004b; Wargo *et al*. 2007b; Huijben *et al*. 2010). In our experiments the AS-selected lines were not set to compete with the reference line, which may explain the absence of costs. We instead obtained an opposite trend, whereby the drug-selected lines (AS2 in particular) fared better in terms of parasitaemia than its unselected counterparts. One possible explanation for these results, which would warrant further studies, is that drug selection may be expected to select for traits that increase the parasite’s survival probabilities in the presence of drugs (Birget et al 2018). Highly replicating parasite lines may be selected for because they can rapidly recover high parasite densities after a bout of drug-induced mortality (Schneider et al. 2008, Schneider et al. 2012). Another possibility is that when confronted with the stressful environment imposed by the drugs, parasites may parasites may exercise a reproductive restraint (Schneider & Reece 2021), that is, sacrifice gametocyte production and prioritize asexual replication to maximise long term survival in the host. Either way, in as much as virulence is related to parasite fitness through high asexual parasite densities and gametocyte production (Mackinnon and Read 2004), this opens the unpalatable possibility of drugs selecting for more resistant but also more virulent parasites. In our experiments, we obtained some indication of a differential virulence between lines. Birds infected with AS1 and AS2 lines had a significantly higher weight loss than those infected with the reference line, but this was not associated to any clear pattern of RBC loss (anaemia). Current evidence for an association between drug resistance alleles and parasite virulence in the field is weak (Tukbasibwe et al. 2017, Cuu et al. 2020), but more work using experimental animal models under controlled conditions are, in our opinion, urgently needed.

Despite its crucial implications for the emergence and spread of drug resistant mutations in the field, work on the transmissibility of artemisinin-resistant parasites, is practically non-existent. (St. Laurent et al. 2015) showed that artemisinin-resistant strains of *P. falciparum* from Cambodia were able to infect and produce sporozoites in three vector species (*An. dirus, An. minimus, An. coluzzi*), and (Pollitt et al. 2014) showed that in the rodent malaria species *P. chabaudi*, the AS-selected line produced more gametocytes (the blood stages of the parasite that are transmitted to the mosquito) than its reference counterpart. Here, we infected mosquitoes with the drug-selected and reference lines in two different experiments. In Experiment 2, all mosquitoes were dissected on a single day (half of the mosquitoes on day 8 and half on day 9). corresponding to the peak ocystaemia as established in previous experiments with the ancestral, reference line In this experiment we observed no significant difference in prevalence between the lines, but a highly significant difference in the number of oocysts between the reference and the two drug-selected lines: the latter produced between 3-5 times fewer oocysts than the reference line (**Figure 3**). This difference could reflect either a true difference in oocyst burden between the lines, or a shift in the within-mosquito dynamics (peak oocystaemia may have occurred earlier or later in the drug-selected lines). To discriminate between these two options a new experiment (Experiment 3) was launched, where both oocysts and sporozoites were quantified at regular, 2-day, intervals, from day 6 to day 20 post infection. This experiment suffered from the caveat of an unusually low infection prevalence (23%), which significantly reduced the statistical power of the burden analyses, and precluded any meaningful analyses of peak oocystaemia. The cumulative frequency of oocysts, however, suggested that oocysts accumulate at a slower rate in mosquitoes infected with the AS-selected lines (**Figure 5**). Further work should explore this issue by, eg measuring potential differences in the growth rate (size) of oocysts issued from reference and AS-resistant lines. The mean overall prevalence and burden of oocysts and sporozoites (i.e. prevalence and burden averaged over all time points) was also in lower AS1 and AS2 mosquitoes. Sporozoite burden, in particular, was 4-5 lower in the drug-selected lines than in the reference line. The results therefore concur with those obtained in the previous experiment in that AS-selected lines were less infective to the mosquitoes than their reference counterparts. Another potential caveat of these experiments was that, for logistical reasons, we only used 2 bird hosts per line. It would be interesting to repeat them with a higher number of hosts to account for potential between-host variability in mosquito infection rates.

In conclusion, our data using drug-selected strains of *P. relictum* provides further proof that susceptibility to artesunate can be rapidly lost under drug pressure. The clearance delay and recrudescence observed during and after the AS treatment, respectively, in two of our three AS-selected lines were similar to the resistance phenotype seen in human malaria infections. The genetic underpinnings of artemisinin resistance are extremely complex and not yet entirely elucidated (Behrens et al. 2021). In *P. falciparum* the resistant phenotype is associated with several mutations in the propeller domain of kelch13 (*pfk13*), a protein that is essential for the intra-erythrocytc growth of the parasite. There are currently 33 candidate *pfk13* mutations associated to artemisinin resistance (WHO Report 2021). In addition, non-synonymous mutations in several other (kelch13-independent) proteins, have also been associated with resistance, either alone or in combination with *pfk13* mutations (Behrens et al. 2021). kelch13 is a highly conserved protein amongst Apicomplexans, and orthologous aminoacid sequences have been found in 21 different *Plasmodium* species, including *P. relictum* (Coppée et al. 2019). The phylogenetic relationship between the k13 in *P. relictum* and human malaria has not been clearly established (Coppée et al. 2019), and its role in the resistance to artesunate in avian malaria remains to be demonstrated.

The recent *de novo* emergence and clonal expansion of a new *pfk13* mutation (R561H) in Subsaharan Africa (Uwimana et al. 2020) where >90% of malaria cases occur, highlights the urgent need to understand the costs associated to artemisinin-resistant strains both in the vertebrate host and in the mosquito vector. In our avian malaria model, we observed no obvious costs of the AS-selected lines in the vertebrate host in single infections. Further work should aim to develop markers of resistance that would allow us to explore whether the drug-selected lines are suppressed when in competition with the reference line (De Roode et al. 2004; Wargo et al. 2007; Huijben et al. 2010). Arguably, however, since our AS-selection lines were not clonal, they may have contained a mixture of resistant and ancestral parasites.

Current knowledge on the transmission potential of artemisin-resistant strains is extremely limited. Our results suggest that, under certain experimental conditions, drug-selected strains may generate lower sporozoite prevalences and burdens in mosquitoes, thus potentially impacting their transmission efficiency. Although the relative costs and benefits of resistance likely differ in magnitude between human and avian malaria parasites, these results provide proof of principle of the need for further studies comparing the infectiousness to mosquitoes of artemisinin-resistant and reference strains of human malaria.

## Acknowledgments

Preprint version 3 of this article has been peer-reviewed and recommended by Peer Community In Evolutionary Biology (https://doi.org/10.24072/pci.evolbiol.100156). We would like to thank the PCI Evolutionary Biology recommender (Sylvie Huijben) and the two reviewers (Sarah Reece and Marianna Szucs) for their thorough review of our manuscript and for their pertinent and useful suggestions. We also thank Bethsabé Sheid, the insectarium (a.k.a Vectopole) coordinator.

## Data, scripts, code, and supplementary information availability

Data available online: https://zenodo.org/record/7113241

## Conflict of interest disclosure

The authors declare that they comply with the PCI rule of having no financial conflicts of interest in relation to the content of the article.

## Funding

The experiments were funded through an ANR-16-CE35-0001-01 (‘EVODRUG’) to Ana Rivero. The Vectopole is one of the platforms of the Vectopole Sud network and is funded through the ANR “Investissements d’avenir” program (ANR-10-LABX-04-01).

## Appendix

## Appendix 1 Supplementary Tables

**ST1:**
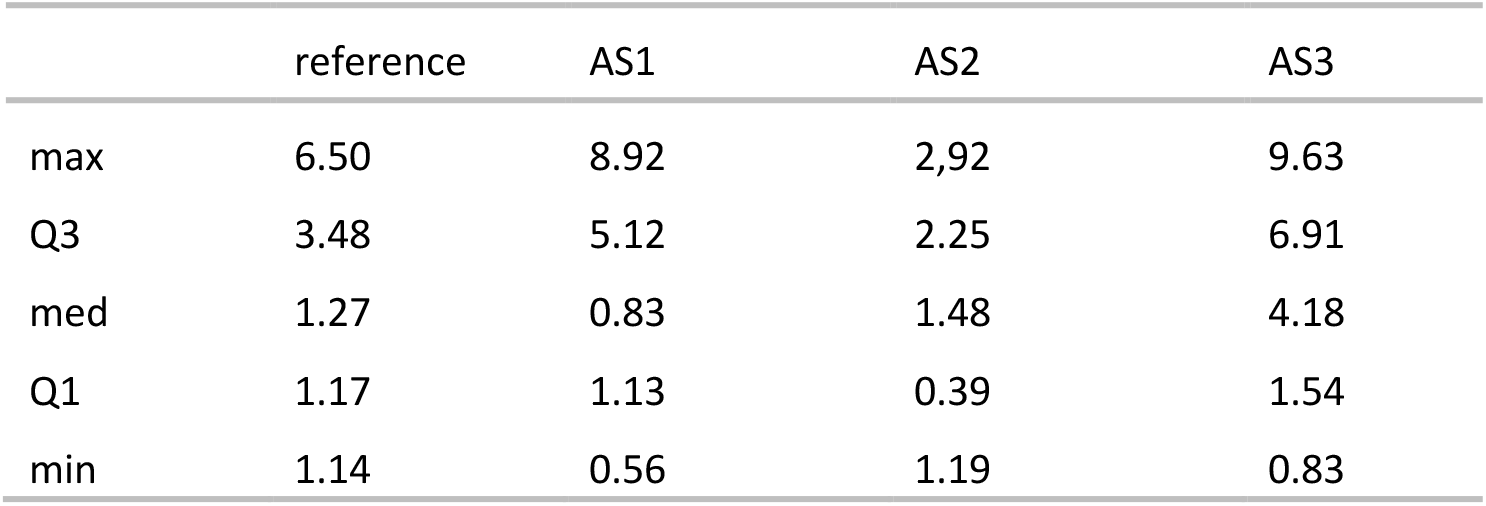
Proportion of blood cells infected (parasitaemia) immediately before the AS-treatment. Table shows the maximum (max) and minimum (min) values, the median (med) and the 25% (Q1) and 75% (Q3) quartiles.

**ST2:**
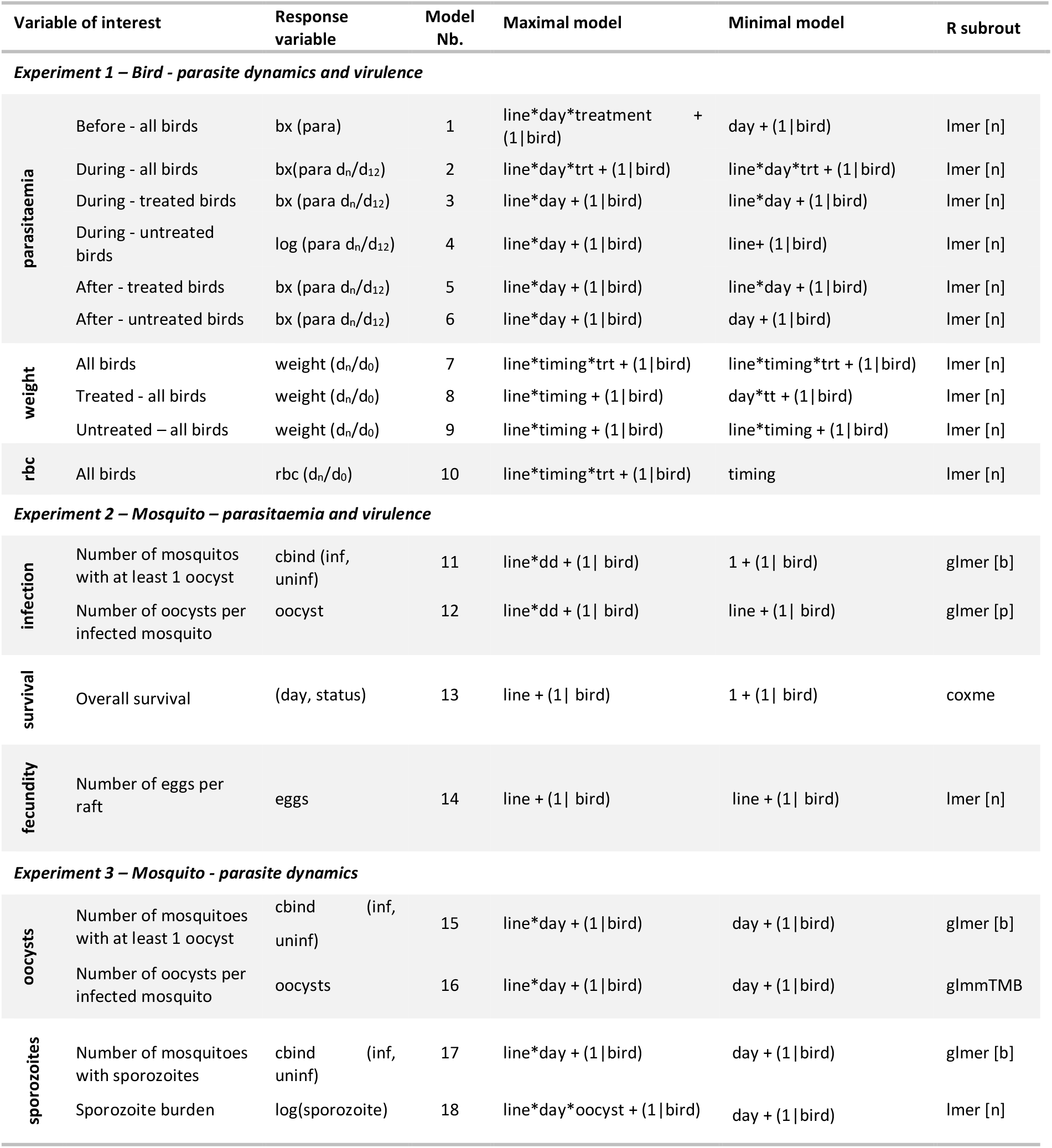
Description of the statistical models used to analyze the costs of AS-resistance in birds and vectors. N gives the number of mosquitoes or birds included in each analysis. “Maximal model” represents the complete set of explanatory variables (and their interactions) included in the model. “Minimal model” represents the model containing only the significant variables and their interactions. N denotes the total number of observations (nb: which is not always the same as the number of independent replicates). Round brackets indicate that the variable was fitted as a random factor. Square brackets indicate the error structure used (n: normal errors, b: binomial errors), *day*: sampling day, *timing:* whether it’s before, during or after the treatment, *status:* alive/dead on sampling day, *eggs:* number of eggs laid, *trt:* treatment (AS or sham-treated), *line:* parasite line, *para:* proportion of infected red blood cells, *rbc*: number of red blood cells per ml.

**ST3:**
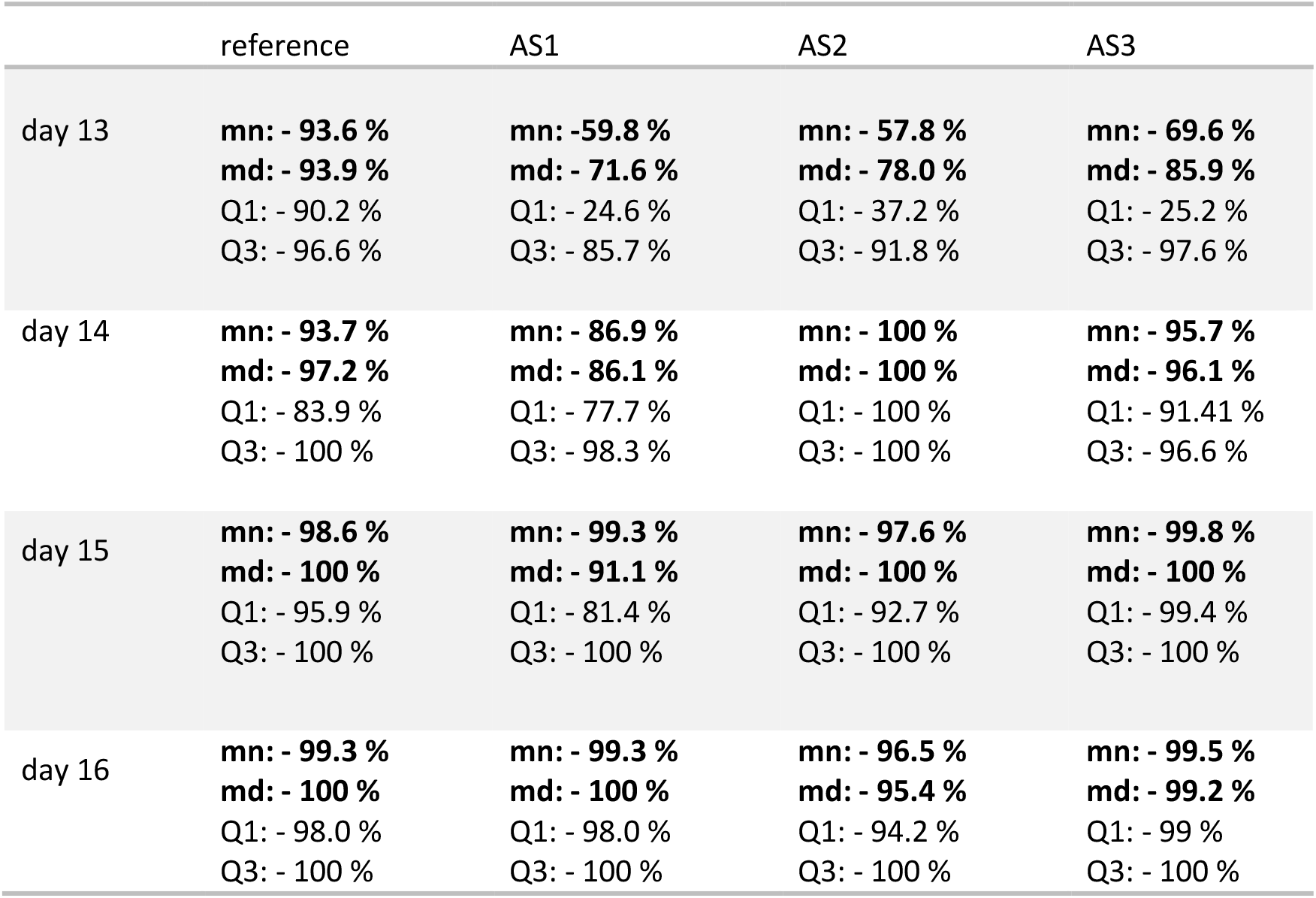
Reduction in parasitaemia as a result of the AS treatment for each of the 3 lines. Treatment was given on days 12-15, parasitaemia was calculated 24h after each treatment (days 13-16). The reduction in parasitaemia is calculated as (*px-pb)/pb * 100* where *px*= parasitaemia at a given day, and *pb* = baseline parasitaemia (parasitaemia immediately before the treatment, day 12). mn=mean, md=median, Q1=25% quartile, Q3= 75% quartile.

**ST4:**
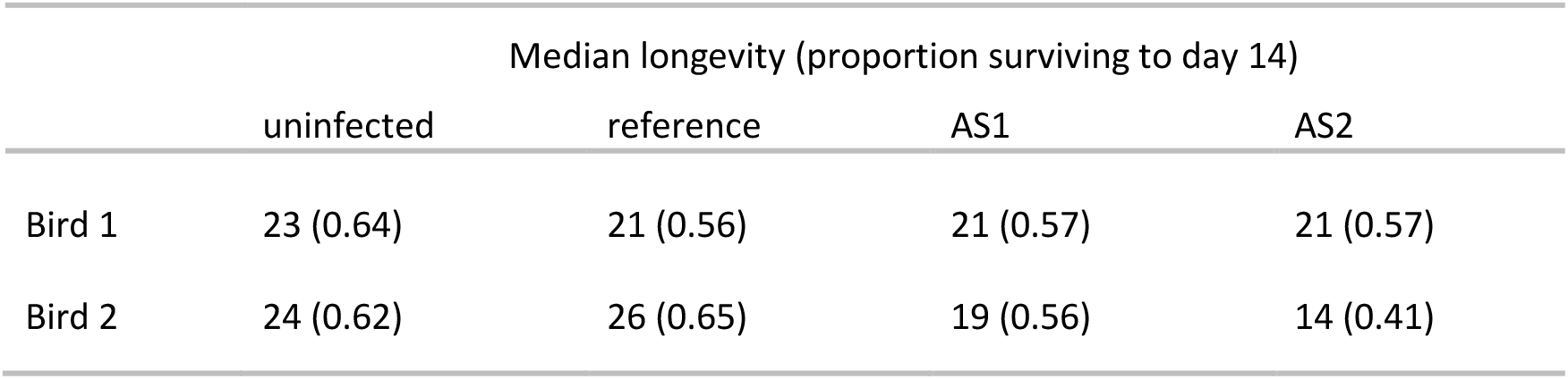
Median mosquito longevity and, in parenthesis, proportion of mosquitoes surviving to day 14 (peak sporozoite production) for uninfected mosquitoes, and mosquitoes infected with the reference or AS-selected lines. Data are presented separately for each bird.

## Appendix 2: Supplementary Figures

**SF1:**
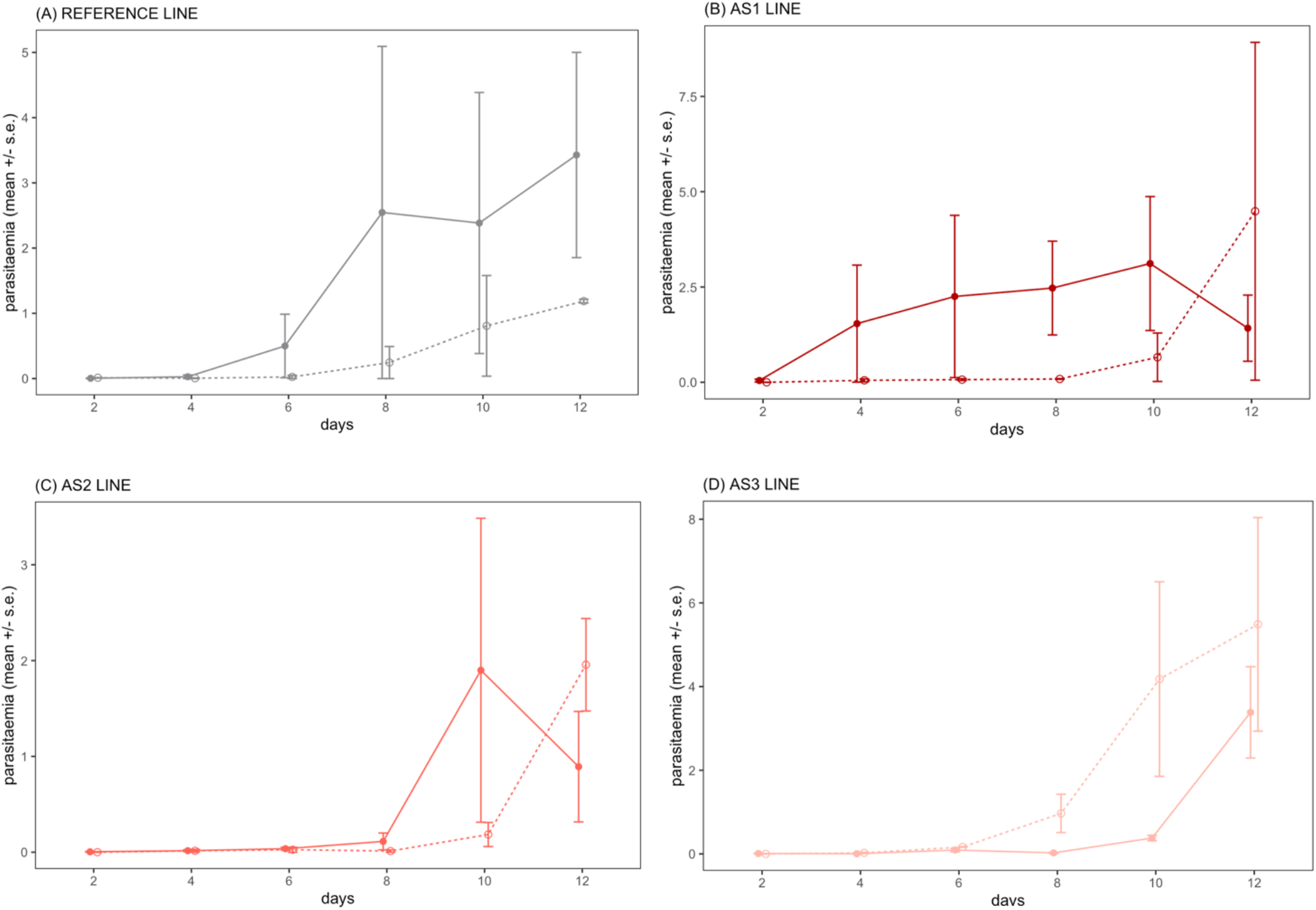
Parasite dynamics before the treatment. (A) reference line, (B) AS1 line, (C) AS2 line, (D) AS3 line. On day 12, birds were blindly allocated to either the untreated (sham injection, dashed lines and empty circles) or treated (artesunate injection, solid lines and full circles) group.

**SF2:**
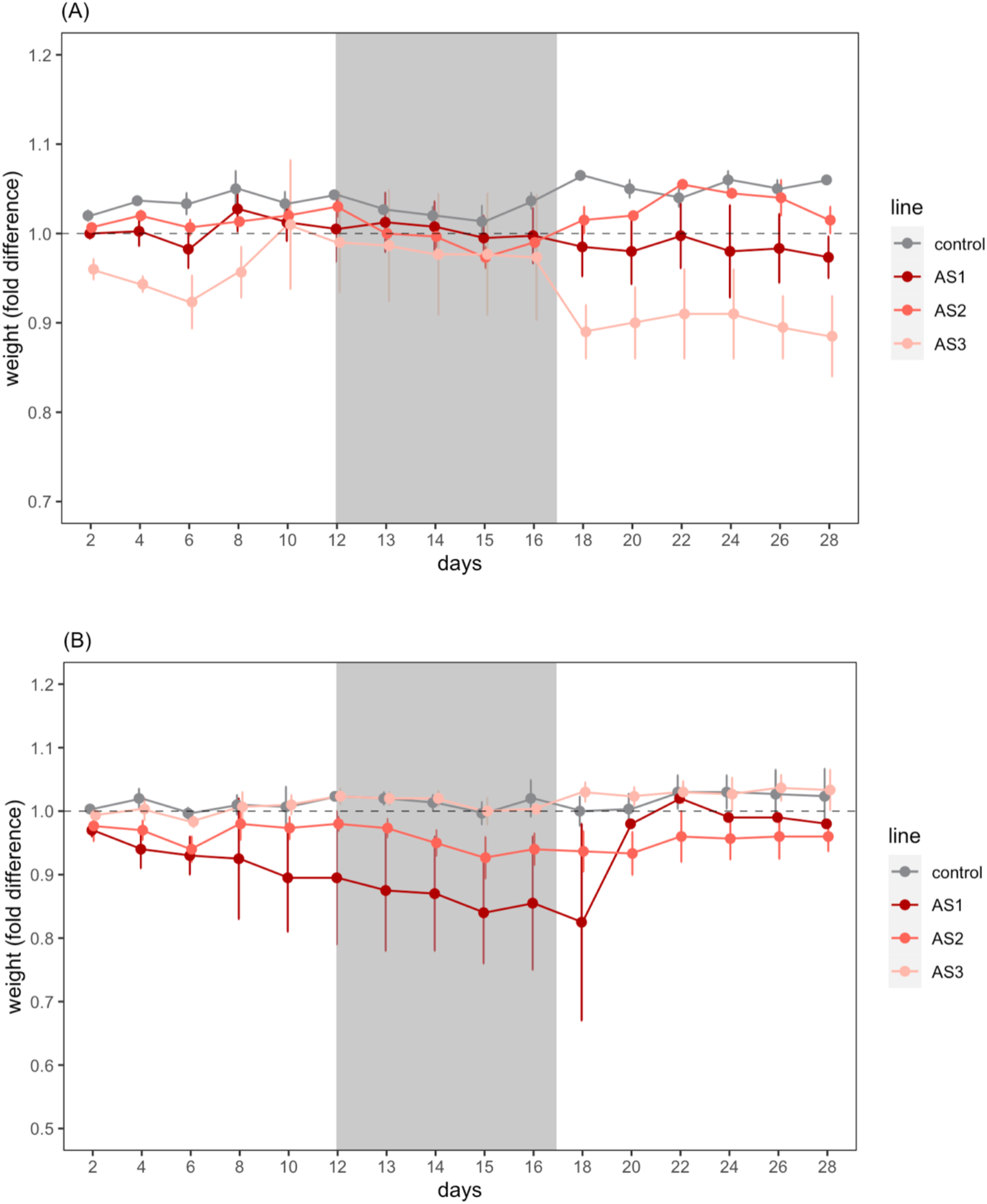
Bird weight changes (mean ± s.e) in artesunate-treated (**A**) and untreated (**B**) birds. Shaded area corresponds to the 4-day treatment period. Untreated birds were sham injected with the artesunate solvent. Dashed line indicates the baseline bird weight at the start of the experiment (day 0). Bars above/below the dashed line indicate an increase/decrease in parasitaemia with respect to day 0.

**SF3:**
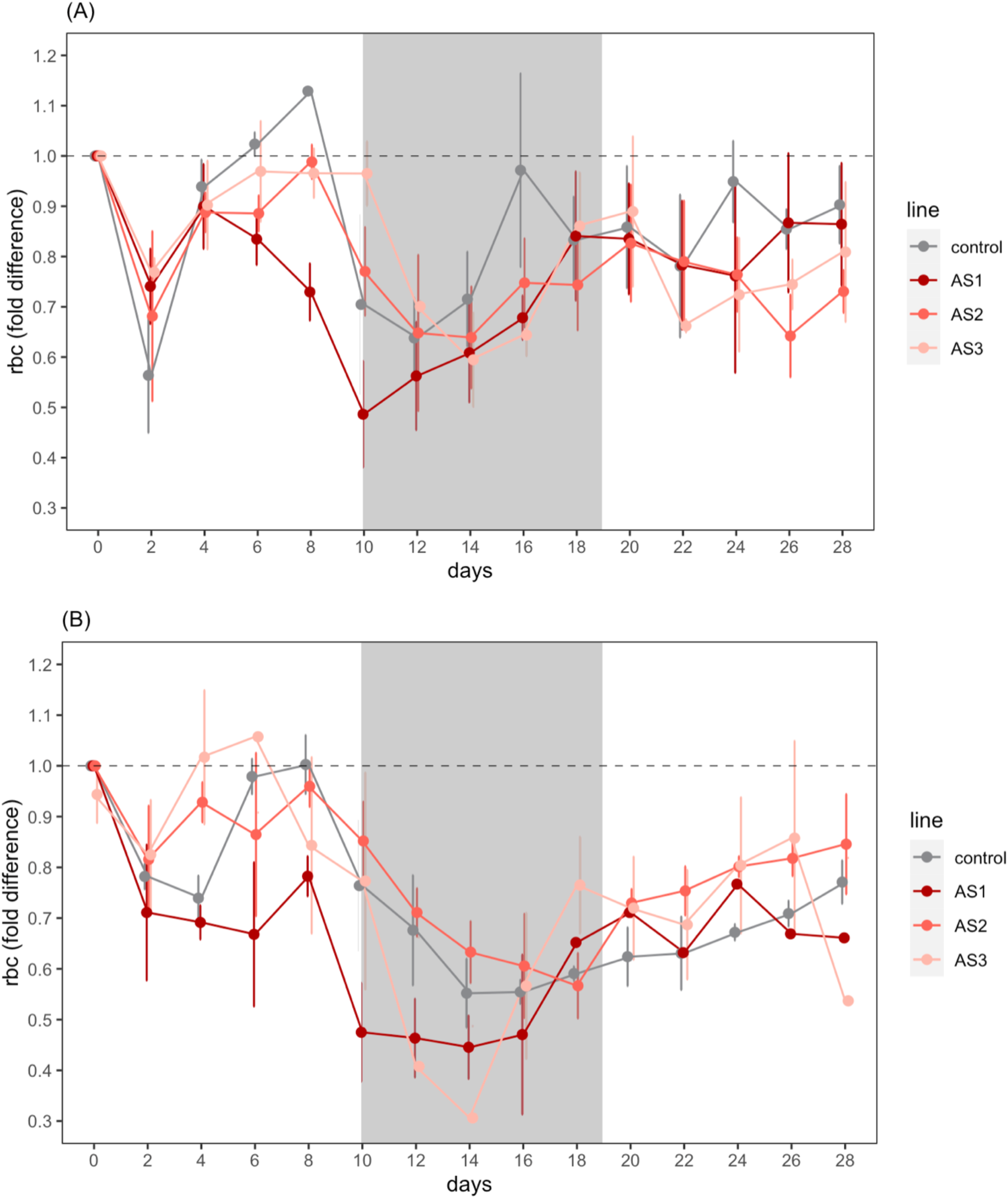
Bird red blood cell count (rbc) changes (mean ± s.e) in artesunate-treated (**A**) and untreated (**B**) birds. Shaded area corresponds to the 4-day treatment period. Untreated birds were sham injected with the artesunate solvent. Dashed line indicates the baseline rbc at the start of the experiment (day 0). Bars above/below the dashed line indicate an increase/decrease in rbc with respect to day 0.

## Notes

### Competing Interest Statement

The authors have declared no competing interest.

### Summary of Updates

Article recommended by PCI Evolutionary Biology Revised to include the comments by the recommenders and two reviewers

