## Supplementary Tables and Figures for "Fitness costs and benefits in response to artificial artesunate selection in *Plasmodium*"

Peer Community In
Evolutionary Biology

RESEARCH ARTICLE

Open Access

Open Data

Open Code

Open Peer-Review

### Fitness costs and benefits in response to artificial artesunate selection in *Plasmodium*

Villa Manon[1], Berthomieu Arnaud[1], Rivero Ana*[1]

[1] MIVEGEC (CNRS, Université de Montpellier, IRD) – Montpellier, France
*Corresponding author

**Cite as**
Villa M. et al. 2022. Fitness costs and benefits in response to artificial artesunate selection in Plasmodium. bioRxiv, 20220128478164, ver 3 peer-reviewed and recommended by Peer Community in Evolutionary Biology. https://doi.org/10.1101/2022.01.28.478164

**Correspondence**


**Recommender**
Silvie Huijben

**Reviewers**
Sarah Reece
Marianna Szucs

This version of the article has been peer-reviewed and recommended by *Peer Community in Evolutionary Biology*
https://doi.org/10.24072/pci.evolbiol.100156

**ABSTRACT**

Drug resistance is a major issue in the control of malaria. Mutations linked to drug resistance often target key metabolic pathways and are therefore expected to be associated with biological costs. The spread of drug resistance depends on the balance between the benefits that these mutations provide in the drug-treated host and the costs they incur in the untreated host. The latter may therefore be expressed both in the vertebrate host and in the vector. Research on the costs of drug resistance focusses on interactions with vertebrate host, yet whether they are also expressed in the vector has been overlooked. In this study, we aim to identify the costs and benefits of resistance against artesunate (AS), one of the main artemisinin derivatives used in malaria-endemic countries. For this purpose, we compared different AS-selected lines of the avian malaria parasite *Plasmodium relictum* to their ancestral (unselected) counterpart. We tested their within host dynamics and virulence both in the vertebrate host and in its natural vector, the mosquito *Culex quinquefasciatus*. The within-host dynamics of the AS-selected lines in the treated birds was consistent with the phenotype of resistance described in human *P. falciparum* malaria: a clearance delay during the treatment followed by a recrudescence once the treatment was interrupted. In the absence of treatment, however, we found no significant costs of resistance in the bird. The results of the two experiments to establish the infectivity of the lines to mosquitoes point towards a decreased infectivity of the drug-selected lines as compared to the ancestral, reference one. We discuss the potential implication of these results on the spread of artesunate resistance in the field.

***Keywords:*** drug-resistance, fitness costs, within-host dynamics, virulence, ACTs, artemisinin derivatives, kelch-13,

### Appendix

**Appendix 1: Supplementary Tables**

| | reference | AS1 | AS2 | AS3 |
|---|---|---|---|---|
| max | 6.50 | 8.92 | 2,92 | 9.63 |
| Q3 | 3.48 | 5.12 | 2.25 | 6.91 |
| med | 1.27 | 0.83 | 1.48 | 4.18 |
| Q1 | 1.17 | 1.13 | 0.39 | 1.54 |
| min | 1.14 | 0.56 | 1.19 | 0.83 |

**ST1:** Proportion of blood cells infected (parasitaemia) immediately before the AS-treatment. Table shows the maximum (max) and minimum (min) values, the median (med) and the 25% (Q1) and 75% (Q3) quartiles.

<table>
<tr><th colspan="2">Variable of interest</th><th>Response variable</th><th>Model Nb.</th><th>Maximal model</th><th>Minimal model</th><th>R subrout</th></tr>
<tr><td colspan="7">Experiment 1 – Bird - parasite dynamics and virulence</td></tr>
<tr><td rowspan="6">parasitaemia</td><td>Before - all birds</td><td>bx (para)</td><td>1</td><td>line*day*treatment + (1|bird)</td><td>day + (1|bird)</td><td>lmer [n]</td></tr>
<tr><td>During - all birds</td><td>bx(para $d_n/d_{12}$)</td><td>2</td><td>line*day*trt + (1|bird)</td><td>line*day*trt + (1|bird)</td><td>lmer [n]</td></tr>
<tr><td>During - treated birds</td><td>bx (para $d_n/d_{12}$)</td><td>3</td><td>line*day + (1|bird)</td><td>line*day + (1|bird)</td><td>lmer [n]</td></tr>
<tr><td>During - untreated birds</td><td>log (para $d_n/d_{12}$)</td><td>4</td><td>line*day + (1|bird)</td><td>line+ (1|bird)</td><td>lmer [n]</td></tr>
<tr><td>After - treated birds</td><td>bx (para $d_n/d_{12}$)</td><td>5</td><td>line*day + (1|bird)</td><td>line*day + (1|bird)</td><td>lmer [n]</td></tr>
<tr><td>After - untreated birds</td><td>bx (para $d_n/d_{12}$)</td><td>6</td><td>line*day + (1|bird)</td><td>day + (1|bird)</td><td>lmer [n]</td></tr>
<tr><td rowspan="3">weight</td><td>All birds</td><td>weight ($d_n/d_0$)</td><td>7</td><td>line*timing*trt + (1|bird)</td><td>line*timing*trt + (1|bird)</td><td>lmer [n]</td></tr>
<tr><td>Treated - all birds</td><td>weight ($d_n/d_0$)</td><td>8</td><td>line*timing + (1|bird)</td><td>day*tt + (1|bird)</td><td>lmer [n]</td></tr>
<tr><td>Untreated – all birds</td><td>weight ($d_n/d_0$)</td><td>9</td><td>line*timing + (1|bird)</td><td>line*timing + (1|bird)</td><td>lmer [n]</td></tr>
<tr><td>rbc</td><td>All birds</td><td>rbc ($d_n/d_0$)</td><td>10</td><td>line*timing*trt + (1|bird)</td><td>timing</td><td>lmer [n]</td></tr>
<tr><td colspan="7">Experiment 2 – Mosquito – parasitaemia and virulence</td></tr>
<tr><td rowspan="2">infection</td><td>Number of mosquitos with at least 1 oocyst</td><td>cbind (inf, uninf)</td><td>11</td><td>line*dd + (1| bird)</td><td>1 + (1| bird)</td><td>glmer [b]</td></tr>
<tr><td>Number of oocysts per infected mosquito</td><td>oocyst</td><td>12</td><td>line*dd + (1| bird)</td><td>line + (1| bird)</td><td>glmer [p]</td></tr>
<tr><td>survival</td><td>Overall survival</td><td>(day, status)</td><td>13</td><td>line + (1| bird)</td><td>1 + (1| bird)</td><td>coxme</td></tr>
<tr><td>fecundity</td><td>Number of eggs per raft</td><td>eggs</td><td>14</td><td>line + (1| bird)</td><td>line + (1| bird)</td><td>lmer [n]</td></tr>
<tr><td colspan="7">Experiment 3 – Mosquito - parasite dynamics</td></tr>
<tr><td rowspan="2">oocysts</td><td>Number of mosquitoes with at least 1 oocyst</td><td>cbind (inf, uninf)</td><td>15</td><td>line*day + (1|bird)</td><td>day + (1|bird)</td><td>glmer [b]</td></tr>
<tr><td>Number of oocysts per infected mosquito</td><td>oocysts</td><td>16</td><td>line*day + (1|bird)</td><td>day + (1|bird)</td><td>glmmTMB</td></tr>
<tr><td rowspan="2">sporozoites</td><td>Number of mosquitoes with sporozoites</td><td>cbind (inf, uninf)</td><td>17</td><td>line*day + (1|bird)</td><td>day + (1|bird)</td><td>glmer [b]</td></tr>
<tr><td>Sporozoite burden</td><td>log(sporozoite)</td><td>18</td><td>line*day*oocyst + (1|bird)</td><td>day + (1|bird)</td><td>lmer [n]</td></tr>
</table>

**ST2:** Description of the statistical models used to analyze the costs of AS-resistance in birds and vectors. N gives the number of mosquitoes or birds included in each analysis. "Maximal model" represents the complete set of explanatory variables (and their interactions) included in the model. "Minimal model" represents the model containing only the significant variables and their interactions. N denotes the total number of observations (nb: which is not always the same as the number of independent replicates). Round brackets indicate that the variable was fitted as a random factor. Square brackets indicate the error structure used (n: normal errors, b: binomial errors), *day*: sampling day, *timing*: whether it's before, during or after the treatment, *status*: alive/dead on sampling day, *eggs*: number of eggs laid, *trt*: treatment (AS or sham-treated), *line*: parasite line, *para*: proportion of infected red blood cells, *rbc*: number of red blood cells per ml.

| | reference | AS1 | AS2 | AS3 |
|---|---|---|---|---|
| day 13 | **mn: - 93.6 %**<br>**md: - 93.9 %**<br>Q1: - 90.2 %<br>Q3: - 96.6 % | **mn: -59.8 %**<br>**md: - 71.6 %**<br>Q1: - 24.6 %<br>Q3: - 85.7 % | **mn: - 57.8 %**<br>**md: - 78.0 %**<br>Q1: - 37.2 %<br>Q3: - 91.8 % | **mn: - 69.6 %**<br>**md: - 85.9 %**<br>Q1: - 25.2 %<br>Q3: - 97.6 % |
| day 14 | **mn: - 93.7 %**<br>**md: - 97.2 %**<br>Q1: - 83.9 %<br>Q3: - 100 % | **mn: - 86.9 %**<br>**md: - 86.1 %**<br>Q1: - 77.7 %<br>Q3: - 98.3 % | **mn: - 100 %**<br>**md: - 100 %**<br>Q1: - 100 %<br>Q3: - 100 % | **mn: - 95.7 %**<br>**md: - 96.1 %**<br>Q1: - 91.41 %<br>Q3: - 96.6 % |
| day 15 | **mn: - 98.6 %**<br>**md: - 100 %**<br>Q1: - 95.9 %<br>Q3: - 100 % | **mn: - 99.3 %**<br>**md: - 91.1 %**<br>Q1: - 81.4 %<br>Q3: - 100 % | **mn: - 97.6 %**<br>**md: - 100 %**<br>Q1: - 92.7 %<br>Q3: - 100 % | **mn: - 99.8 %**<br>**md: - 100 %**<br>Q1: - 99.4 %<br>Q3: - 100 % |
| day 16 | **mn: - 99.3 %**<br>**md: - 100 %**<br>Q1: - 98.0 %<br>Q3: - 100 % | **mn: - 99.3 %**<br>**md: - 100 %**<br>Q1: - 98.0 %<br>Q3: - 100 % | **mn: - 96.5 %**<br>**md: - 95.4 %**<br>Q1: - 94.2 %<br>Q3: - 100 % | **mn: - 99.5 %**<br>**md: - 99.2 %**<br>Q1: - 99 %<br>Q3: - 100 % |

**ST3:** Reduction in parasitaemia as a result of the AS treatment for each of the 3 lines. Treatment was given on days 12-15, parasitaemia was calculated 24h after each treatment (days 13-16). The reduction in parasitaemia is calculated as *(px-pb)/pb * 100* where *px*= parasitaemia at a given day, and *pb* = baseline parasitaemia (parasitaemia immediately before the treatment, day 12). mn=mean, md=median, Q1=25% quartile, Q3= 75% quartile.

| | Median longevity (proportion surviving to day 14) | | | |
|---|---|---|---|---|
| | uninfected | reference | AS1 | AS2 |
| Bird 1 | 23 (0.64) | 21 (0.56) | 21 (0.57) | 21 (0.57) |
| Bird 2 | 24 (0.62) | 26 (0.65) | 19 (0.56) | 14 (0.41) |

**ST4:** Median mosquito longevity and, in parenthesis, proportion of mosquitoes surviving to day 14 (peak sporozoite production) for uninfected mosquitoes, and mosquitoes infected with the reference or AS-selected lines. Data are presented separately for each bird.

**Appendix 2: Supplementary Figures**

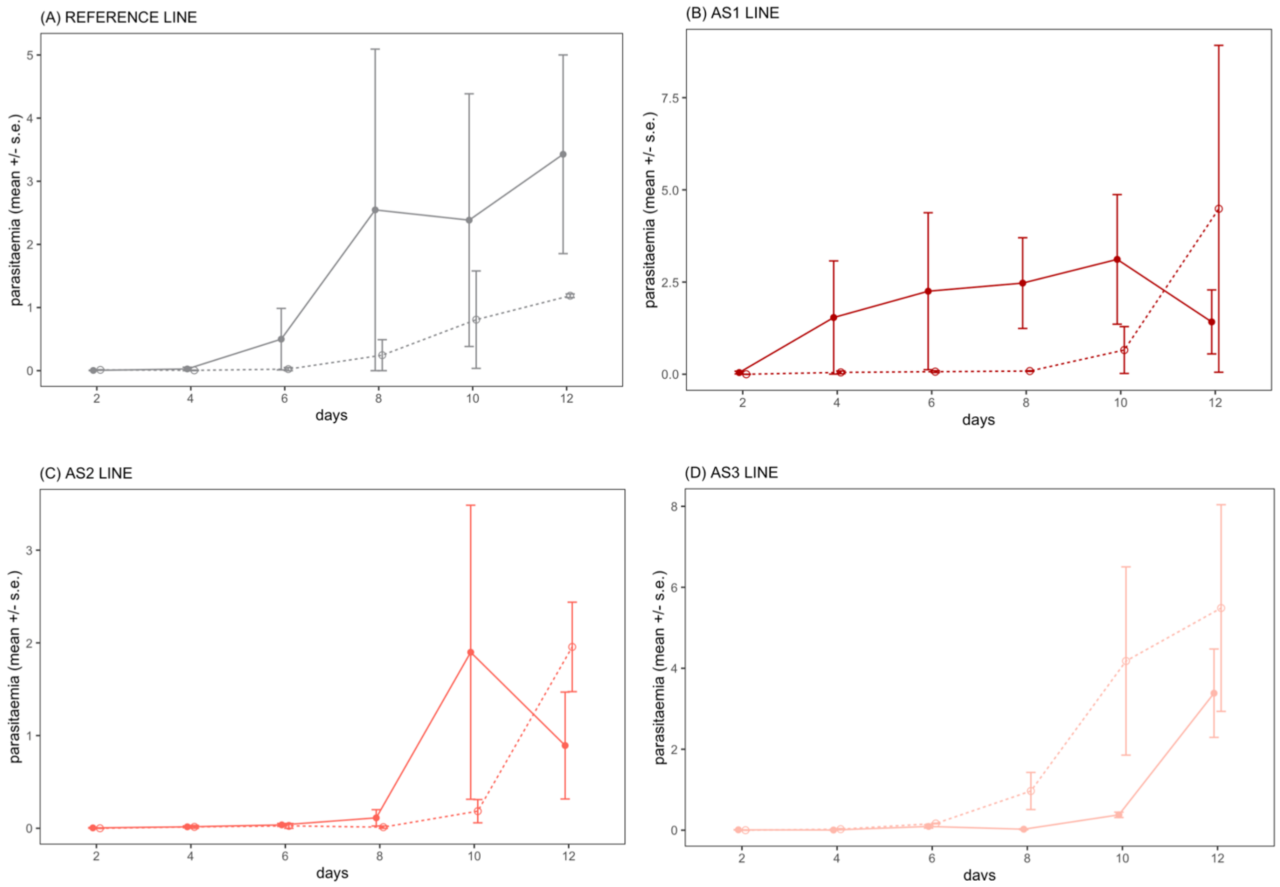


**SF1:** Parasite dynamics before the treatment. (A) reference line, (B) AS1 line, (C) AS2 line, (D) AS3 line. On day 12, birds were blindly allocated to either the untreated (sham injection, dashed lines and empty circles) or treated (artesunate injection, solid lines and full circles) group.

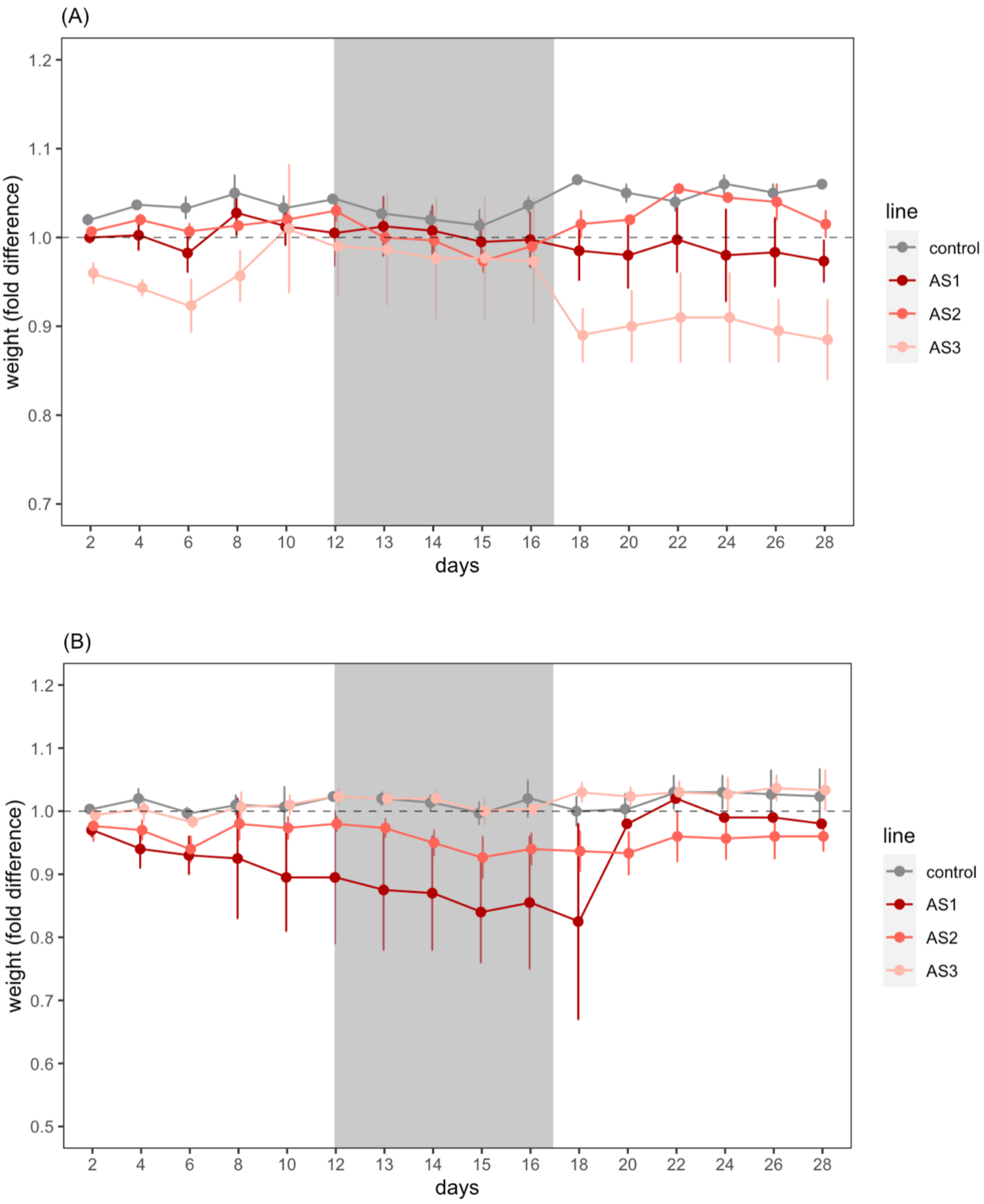


**SF2:** Bird weight changes (mean ± s.e) in artesunate-treated (**A**) and untreated (**B**) birds. Shaded area corresponds to the 4-day treatment period. Untreated birds were sham injected with the artesunate solvent. Dashed line indicates the baseline bird weight at the start of the experiment (day 0). Bars above/below the dashed line indicate an increase/decrease in parasitaemia with respect to day 0.

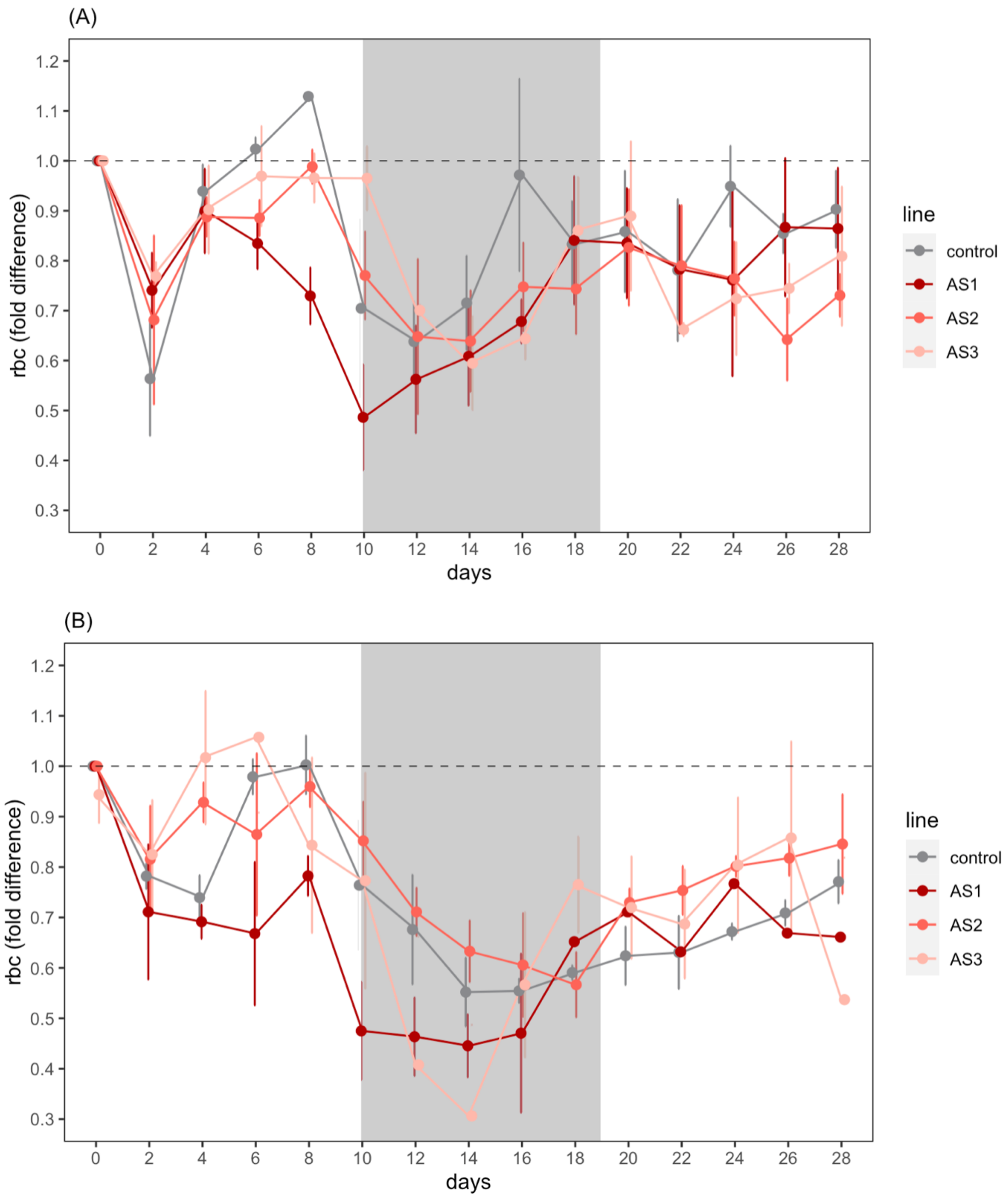


**SF3:** Bird red blood cell count (rbc) changes (mean ± s.e) in artesunate-treated (**A**) and untreated (**B**) birds. Shaded area corresponds to the 4-day treatment period. Untreated birds were sham injected with the artesunate solvent. Dashed line indicates the baseline rbc at the start of the experiment (day 0). Bars above/below the dashed line indicate an increase/decrease in rbc with respect to day 0.
